# The number of anal fin-rays is decided by two heritable traits, anteroposterior length of the anal fin and interval between the anal fin-rays

**DOI:** 10.64898/2025.12.08.692906

**Authors:** Tetsuaki Kimura

## Abstract

The number of median fin-rays is a readily quantifiable trait that has been previously studied due to its interspecific variation. I observed in F2 progeny from two inbred lines of medaka with differing anal fin-ray numbers (Kaga and Hd-rRII1) that individuals exhibited the same ray number but differed in the anteroposterior length of the anal fin and the interval between the anal fin-rays. Inducing vertebral fusion to shorten the anal fin anteroposterior length resulted in a decrease in ray number. Given reports of an increased ray number during low-temperature development, I reared Hd-rRII1 fry at 17°C post-hatching, which resulted in an increase in the number of rays. At this temperature, the anal fin anteroposterior length and the interval between the anal fin-rays were also slightly reduced. Since the posterior boundary of the anal fin is determined by Hox genes, I hypothesize that a temperature-sensitive signaling pathway exists downstream of these genes. Collectively, our results suggest that the medaka anal fin-ray number is determined by at least two genetic traits: the anal fin anteroposterior length and the interval between the anal fin-rays.

## Introduction

The median fins of teleosts have a characteristic exoskeleton structure (fin-ray) (1). The number of anal fin-rays is variable within fish, and is one of the criteria used in fish taxonomy. The ease with which the number of fin rays can be counted has facilitated their genetic analysis (2). In particular, variation in the number of anal fin-rays is a difference first recognized between southern and northern Japanese populations of medaka; it is known to be a heritable trait (3–6). However, the genes that determine the number of anal fin-rays have not been identified to date.

## Results

To identify the genes underlying the number of median fin-rays, I crossed an inbred line from a southern Japanese population, Hd-rRII1, with a northern Japanese population inbred line, Kaga. These inbred lines differed by an average of four fin rays in the anal fin and one fin ray in the dorsal fin (Table 1). The F1 fish had 18.8 rays on average; the number ranged from 18 to 20 (Table 1). Since the two inbred lines and F1 fish possess identical genetic information, the number of anal fin-rays must be decided by both genetic and environmental factors. This is consistent with previous observations (6, 7).

**Table 1.** Number of anal and dorsal fin-rays in each generation.

| Strain | Number of anal fin-rays | | | | | | | | | | Mean ± SEM | Number of dorsal fin-rays | | | Mean ± SEM | Total |
| --- | --- | --- | --- | --- | --- | --- | --- | --- | --- | --- | --- | --- | --- | --- | --- | --- |
|  | 14 | 15 | 16 | 17 | 18 | 19 | 20 | 21 | 22 | 23 |  | 4 | 5 | 6 |  |  |
| Hd-rRII1 | — | — | — | — | 5 | 24 | 6 | — | — | — | 19.0 ± 0.10 | — | 1 | 34 | 5.95 ± 0.03 | 35 |
| Kaga | 4 | 18 | 14 | — | — | — | — | — | — | — | 15.3 ± 0.11 | — | 34 | 2 | 5.05 ± 0.04 | 36 |
| F<sub>1</sub> | — | — | — | — | 15 | 42 | 1 | — | — | — | 18.8 ± 0.06 | — | 2 | 56 | 5.97 ± 0.02 | 58 |
| F<sub>2</sub> | — | 1 | 11 | 52 | 67 | 40 | 26 | 2 | — | 1 | 18.1 ± 0.09 | 2 | 57 | 141 | 5.70 ± 0.03 | 200 |

SEM is standard error of the mean. — indicates that no observation was made.

The F2 fish derived from the F1 intercross had an average of 18.1 anal fin-rays; the number ranged from 15 to 23 (Table 1). I considered that the difference in the number of rays was derived from the anteroposterior length of the anal fin, because the ratio of tail to abdomen differed between the Kaga and Hd-rRII1 strains (8). Since the Kaga and Hd-rRII1 strains differ in terms of vertebrae (8), I superimposed two images of F2 fish with 29 vertebrae. As expected, F2 fish with 19 rays had a more posteriorly elongated anal fin than F2 fish with 17 rays (Fig. 1A). Surprisingly, I also found that the anal fin of some F2 fish with 19 rays was the same length as that of F2 fish with 17 rays (Fig. 1B). This implied that F2 fish showed a variation in the interval between rays. These findings suggested that the number of anal fin-rays was determined by the anteroposterior length of the anal fin and the intervals between rays. Additionally, the pattern of the rays was independent of the segmental pattern of the somite and vertebrae, although the somitic mesoderm contributed to the development of the rays (9).

**Figure 1.**
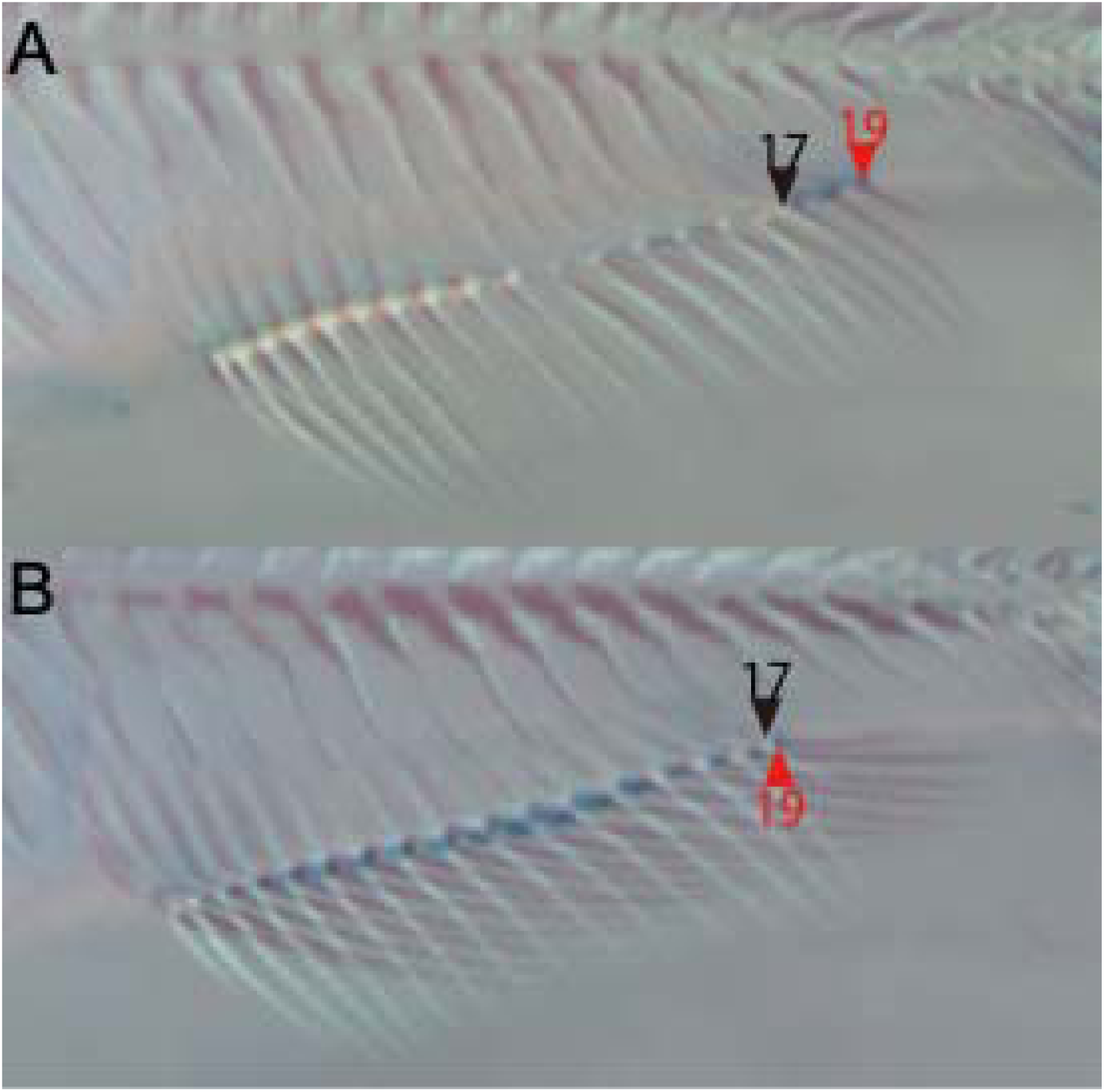
Superimposition of F_2_ anal fin-rays. (A) superimposed image 17 rays and 19 rays F_2_ fish. The 19-ray fish has a longer anal fin than the 17-ray fish. (B) superimposed image 17 rays and another 19 rays F_2_ fish. White 17-ray fish is the same in A. The 19-ray fish has the same length anal fin as the 17-ray fish. This indicates that the 19-ray fish has a narrower interval of fin-rays than the 17-ray fish. All three fish have 29 vertebrae. Note the pattern of vertebrae is the same for both A and B. Arrowheads indicate the most posterior ray. Numbers indicate the total number of fin-rays.

To confirm the hypothesis, I shortened the medaka using a phenylthiourea treatment. Yamamoto and colleagues (10) have previously shown that phenylthiourea treatment induces a nonheritable shortened body length due to vertebral fusion resulting from a twisted notochord, and decreases the number of anal fin-rays. I confirmed that this treatment decreased the number of anal fin-rays, but did not affect the somite segmentation pattern, using Hd-rRII1 (Fig. 2 and Table 2). If the pattern of anal fin-rays was dependent on the pattern of the vertebrae, then the rays were expected to fuse at the position of vertebral fusion. On the other hand, if the pattern of anal fin-rays was dependent on the somite pattern, then the number of anal fin-rays should remain the same while the spacing between rays became narrower. However, neither of these possibilities was found in our results. This again suggested that the number of anal fin-rays is determined by the anteroposterior length of the anal fin, and that the pattern of anal fin-rays is independent of the segmentation pattern of the vertebrae and somite.

**Figure 2.**
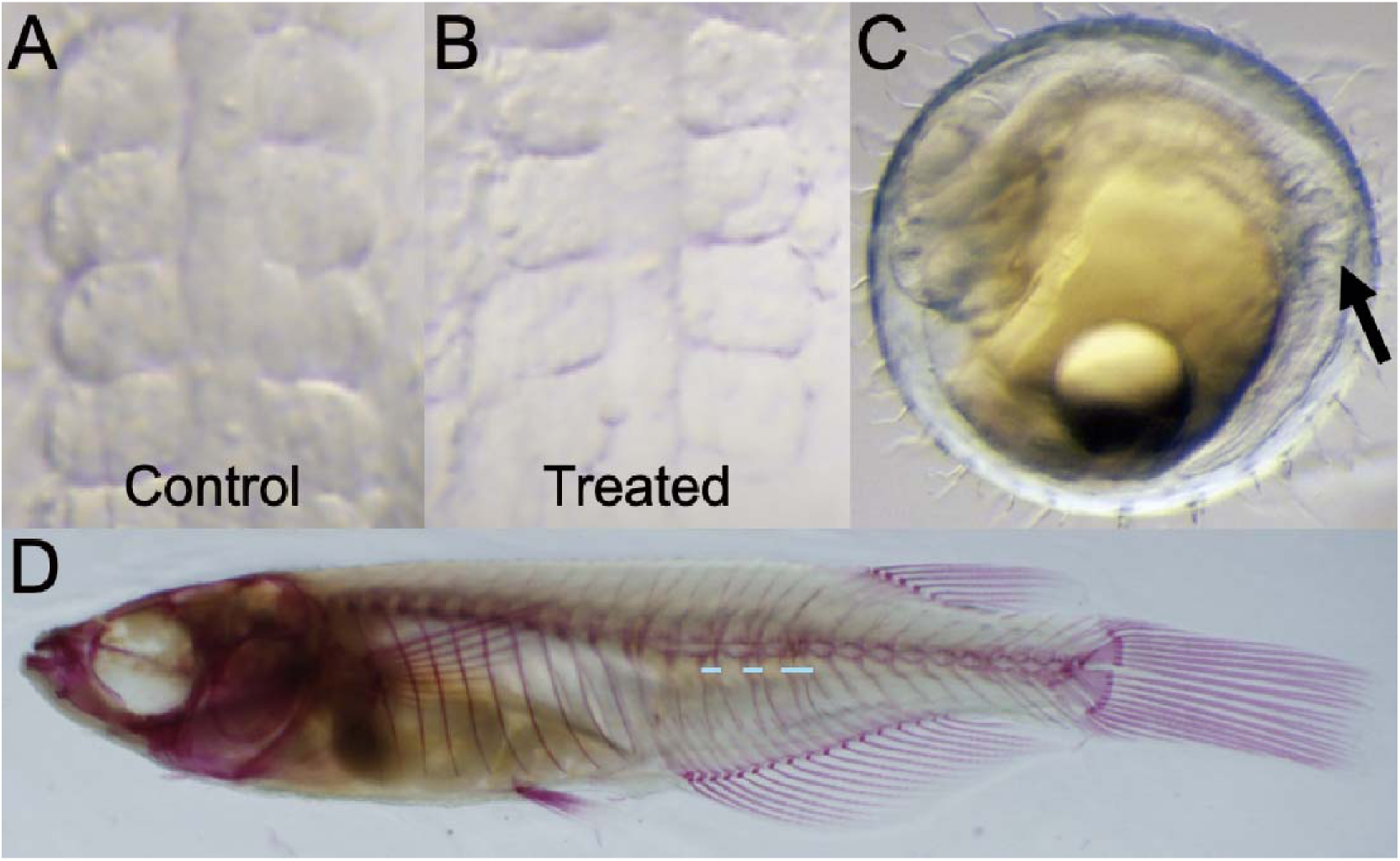
Phenylthiourea treatment induces truncation of body length. (A, B) Segmental pattern of somite in stage 22 embryos. The phenylthiourea treatment does not affect the segmental pattern of somite in A or B. (C) One of the most severe phenotypes at stage 32. The treatment induces notochord vending at a late developmental stage. The arrow indicates the vending notochord. (D) Phenotype of the adult stage. Some vertebral segments are fused (light blue lines). As a result, body length becomes shorter and the number of anal fin-rays decreases to 17 (Hd-rRII1 usually has 19 rays; Table 2). Note the pattern of anal fin-rays is normal, though the vertebral pattern is perturbed.

**Table 2.**
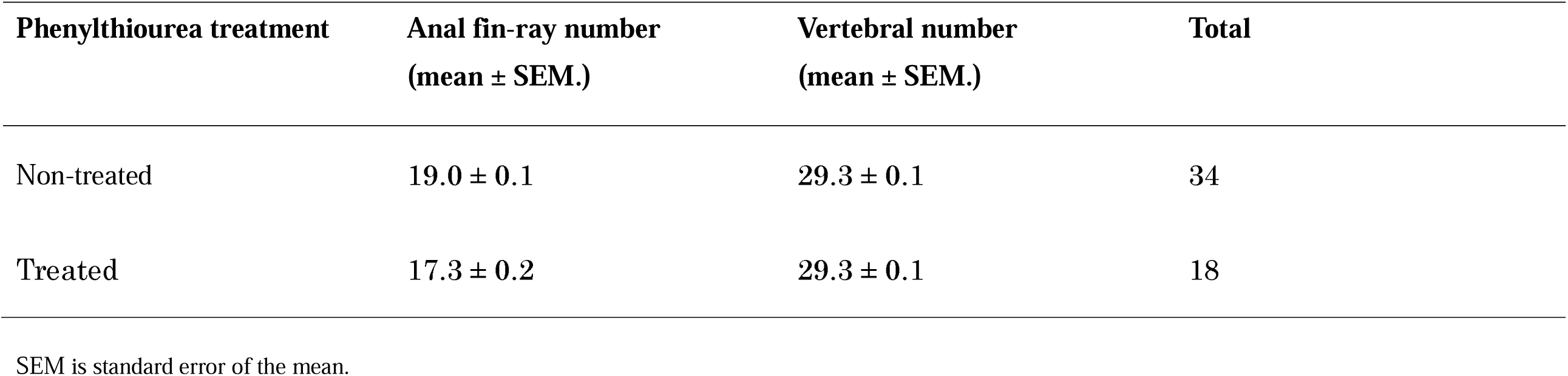
Effect of phenylthiourea treatment.

Reports indicate that rearing at lower temperatures increases anal fin ray number in medaka (6). Given that the posterior boundary of the anal fin is specified by Hox genes (11), I hypothesized that this increase results from a reduction in the spacing between fin rays. To confirm this, Hd-rRII1 fry were reared at two temperatures: 26°C (control) and 17°C (lower temperature) post-hatching. As expected, fish reared at 17°C exhibited a significant increase in fin-ray number (Welch’s *t-test P* = 3.92 × 10^−4^) (Fig. 3B and Table 3), and the spacing between fin-rays was also reduced (Welch’s *t-test P* = 3.97 × 10^−12^) (Fig. 3C and Table 3). Surprisingly, I also observed a reduction in the anteroposterior length of the anal fin itself (Welch’s *t-test P* = 8.23 × 10^−10^) (Fig. 3D and Table 3), which limited the magnitude of the increase in ray number. This temperature perturbation experiment further supports the hypothesis that anal fin-ray number is determined by the anteroposterior length of the anal fin and the spacing between the fin-rays.

**Figure 3.**
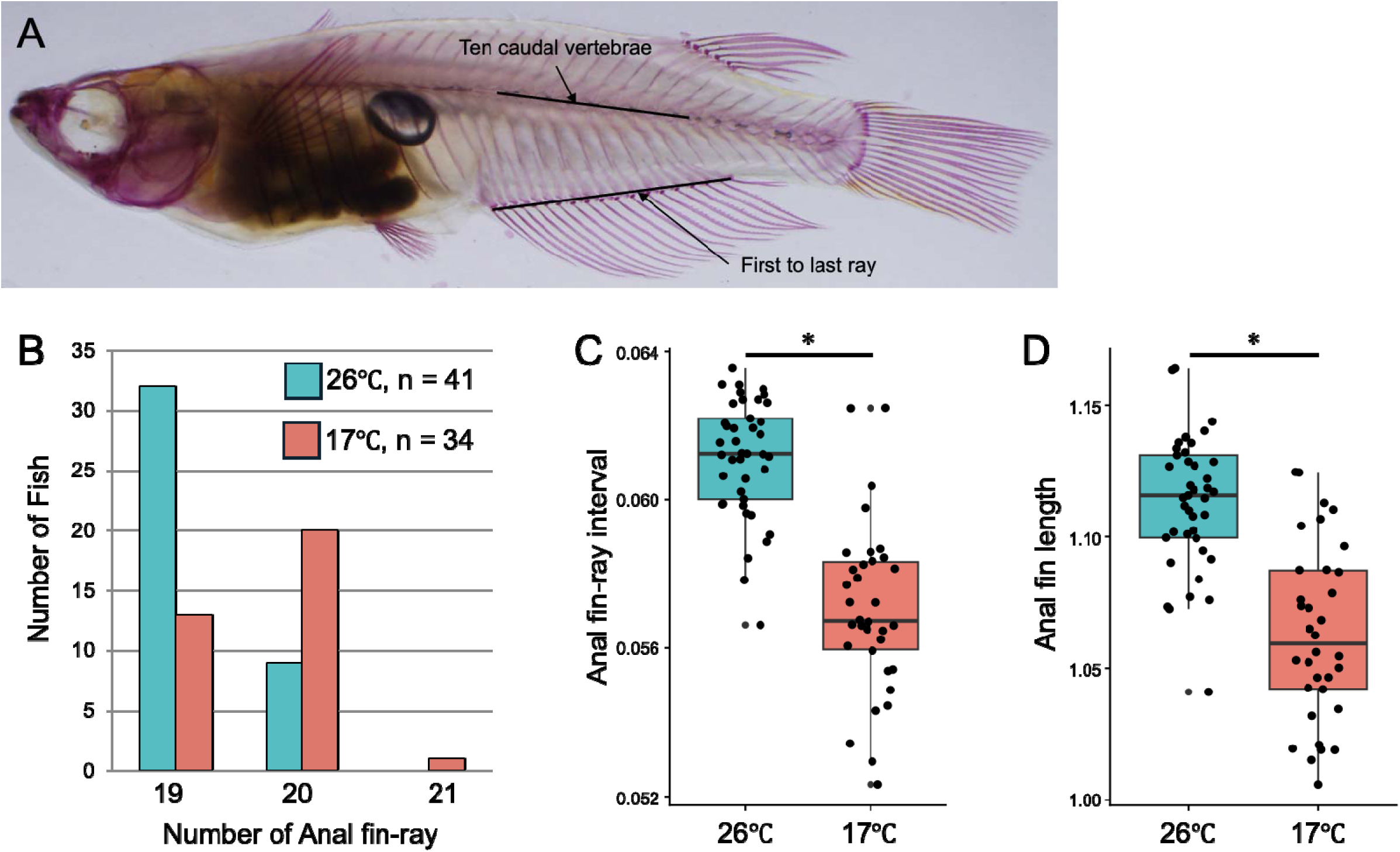
Rearing temperature changes the interval between rays and the anteroposterior anal fin length. (A) Anal fin traits. The anteroposterior length of the anal fin is the ratio of the anteroposterior anal fin length to the length of 10 caudal vertebrae. The interval of the anal fin-rays is the mean of the ray intervals in the anal fin, which is calculated as the anteroposterior length of the anal fin divided by the number of ray intervals. (B) The number of anal fin-rays. Fish reared at 17°C have a larger number of rays. (C) The boxplot shows the anal fin-ray intervals. The thick black lines in the boxes indicate the medians of the groups. The two hinges of the boxes are the first and third quartiles, and the whiskers show the largest/smallest observations that fall within a limit of 1.5 times the box size from the nearest hinge. Fish reared at 17°C have a narrower interval. (D) Anal fin length is also affected by the rearing temperature. The convention of the boxes is the same as in C. All traits show a significant difference according to Welch’s t-test (P < 0.05). Asterisks indicate a significant difference. N indicates the number of fish.

**Table 3.** Rearing temperature affect anal fin traits.

| **Rearing** | **Number of anal fin rays** | **Anal fin ray interval** | **Anal fin length** | **Number of fish** |
| --- | --- | --- | --- | --- |
| **temperature** | **(mean ± SEM.)** | **(mean ± SEM.)** | **(mean ± SEM.)** |  |
| 26°C | 19.2 ± 0.065 | 0.061 ± 0.0002 | 1.11 ± 0.004 | 41 |
| 17°C | 19.6 ± 0.093 | 0.057 ± 0.0004 | 1.06 ± 0.006 | 34 |

SEM is standard error of the mean.

## Discussion

The number of dorsal and anal fin rays was not constant in genetically homogenous inbred lines and their F1 hybrids. This suggests that the ray number is determined by environmental factors, as previously demonstrated (6, 7). However, clear differences in dorsal and anal fin ray numbers were observed between the Kaga and Hd-rRII1 strains, indicating a genetic basis for determining ray number. Analysis of vertebral fusion induction and temperature variation did not reveal statistically significant differences in dorsal fin ray number, likely due to the short postaxial length of the dorsal fin. While statistical significance was not achieved, the average number of dorsal fin rays in fish raised at 17°C tended to be lower than those raised at 26°C (17°C mean ± standard error of the mean (SEM): 5.88 ± 0.056, 26°C mean ± SEM: 5.95 ± 0.034).

The reduction in anal fin ray number following vertebral fusion suggests that median fin rays are newly formed structures independent of the body segment and vertebral pattern. These median fin rays appear to inherit patterns from underlying pterygiophores and develop in an anteroposterior manner. In adult medaka, the spacing between fin rays widens posteriorly. Furthermore, fusions of anterior supporting bones and fin rays are frequently observed, suggesting that this unequal spacing develops during growth. Therefore, genetic analysis of fin ray spacing should ideally measure the intervals between pterygiophores at an early developmental stage (likely when three pterygiophores have formed).

Kaga exhibits fewer dorsal and anal fin rays compared to Hd-rRII1, likely due to its wider fin ray spacing. This phenotype is potentially manifested under the rearing temperature of 26°C. As fin ray spacing decreases at lower temperatures, I hypothesize that Kaga is adapted for survival in colder environments compared to Hd-rRII1, resulting in wider spacing at the same temperature. Given that fin ray spacing relates to the physical strength of the fin, it is expected to converge towards an optimal interval. Experiments involving temperature variation suggest the existence of a morphogen that signals fin ray spacing through a relay or diffusion mechanism. Genetic polymorphisms altering the rate of morphogen diffusion likely underlie the observed variation in spacing between these two inbred lines.

Unexpectedly, the anteroposterior length of the anal fin was reduced under 17°C rearing conditions. This suggests that a morphogen downstream of Hox genes is acting on fin development. However, as I did not confirm the expression domains of Hox genes in this study, a change in Hox expression cannot be ruled out. Future research should investigate whether temperature variation alters the position of Hox gene expression. Furthermore, identifying the morphogen(s) that pattern the underlying pterygiophores is necessary.

## Methods

### Fish

Hd-rR-II1 and Kaga strains (12) are inbred medaka strains established from a southern and a northern Japanese population, respectively (13). These strains were obtained from the National BioResource Project (NBRP) Medaka (www.shigen.nig.ac.jp/medaka/). The medaka were reared at 26.0°C on a 14-hour light/10-hour dark cycle. I incubated all eggs at 28 °C until hatching to minimize the relative effects of temperature. Two pairs of Hd-rR-II1 and Kaga strains were crossed to generate the F1 progeny. Sex combination was reciprocal. A total of 200 F2 progeny were obtained from intercrossing 11 pairs of the F1 progeny.

### Vertebral fusion induction

Induction of vertebral fusion was performed based on the methods of Yamamoto et al. using fertilized eggs from the Hd-rRII1 strain (10).

### Analysis of temperature effects

To investigate the effect of rearing temperature on fin ray number, fertilized eggs were collected from four pairs of Hd-rRII1 fish. Larvae hatched and were raised at either 26°C or 16-17°C for four months. Individuals exhibiting vertebral fusion were frequently observed in the 17°C group and were excluded from analysis.

### Skeletal preparation and phenotype measurement

Skeletal specimens were prepared and phenotypes measured according to the methods of Kimura et al. (8). Due to continuous growth throughout the medaka lifespan, anteroposterior length of the anal fin was calculated as the ratio of the straight-line distance from the first to last fin ray to a length encompassing the ten caudal vertebrae (Fig. 3A). The interval between the anal fin-rays was calculated by dividing the anteroposterior length of the anal fin by the number of ray intervals.

### Statistical analysis

Statistical analyses of each trait were performed using Welch’s t-test in R. A *P* value of 0.05 or less was considered statistically significant.

## Acknowledgments

I thank the National BioResource Project Medaka (NBRP Medaka) of the Ministry of Education, Culture, Sports, Science and Technology (MEXT), Japan for providing Hd-rRII1, Kaga. This work was supported by JSPS KAKENHI Grant Number 24770063.

## References

1. P. M. Mabee, P. L. Crotwell, N. C. Bird, A. C. Burke, Evolution of median fin modules in the axial skeleton of fishes. J Exp Zool. 294, 77–90 (2002).

2. K. M. Nichols, P. A. Wheeler, G. H. Thorgaard, Quantitative trait loci analyses for meristic traits in *Oncorhynchus mykiss*. Environ Biol Fishes. 69, 317–331 (2004).

3. N. Egami, Studies on the variation of the number of the anal fin-rays in *Oryzias latipes* I. Geographical variation in wild populations. Jpn J Ichthyol. 3, 33–35, 87-89 (1953).

4. N. Egami, Studies on the variation of the number of the anal fin-rays in *Oryzias latipes* II. Cross experiments. Jpn J Ichthyol. 3, 171–178 (1954).

5. N. Egami, M. Yoshino, Studies on the variation of the number of the anal fin-rays in *Oryzias latipes* III. Supplementary note on the geographical variation in the wild populations. Jpn J Ichthyol. 7, 83–88 (1958).

6. M. Y. Ali, C. C. Lindsey, Heritable and temperature-induced meristic variation in the medaka, *Oryzias latipes*. Can J Zool. 52, 959–976 (1974).

7. C. C. Lindsey, R. W. Harrington Jr., Extreme vertebral variation induced by temperature in a homozygous clone of the self-fertilizing cyprinodontid fish *Rivulus marmoratus*. Can J Zool. 50, 733–744 (1972).

8. T. Kimura, M. Shinya, K. Naruse, Genetic analysis of vertebral regionalization and number in medaka (*Oryzias latipes*) inbred lines. G3 (Bethesda). 2, 1317–1323 (2012).

9. A. Shimada et al., Trunk exoskeleton in teleosts is mesodermal in origin. Nat Commun. 4, 1639 (2013).

10. T. Yamamoto, H. Tomita, N. Matsuda, Hereditary and non-heritable vertebral anchylosis in medaka, *Oryzias latipes*. Jpn J Genet. 38, 36–47 (1963).

11. R. Koita et al., The phenotypic variation of widefins medaka is due to the insertion of a giant transposon containing a viral genome within hoxca cluster. Genetics. iyaf218 (2025).

12. Hyodo-Taguchi Y., Inbred strains of the medaka, *Oryzias latipes*. Fish Biol. J. Medaka 8: 11–14 (1996).

13. Y. Takehana, N. Nagai, M. Matsuda, K. Tsuchiya, M. Sakaizumi, Geographic variation and diversity of the cytochrome b gene in Japanese wild populations of medaka, Oryzias latipes. Zool Sci. 20, 1279–1291 (2003).

